# cageminer: an R/Bioconductor package to prioritize candidate genes by integrating GWAS and gene coexpression networks

**DOI:** 10.1101/2021.08.04.455037

**Authors:** Fabricio Almeida-Silva, Thiago M. Venancio

**Author notes:** Laboratório de Química e Função de Proteínas e Peptídeos, Centro de Biociências e Biotecnologia, Universidade Estadual do Norte Fluminense Darcy Ribeiro. Av. Alberto Lamego 2000, P5, sala 217, Campos dos Goytacazes, RJ, Brazil.

## Abstract

**Summary:** Although genome-wide association studies (GWAS) identify variants associated with traits of interest, they often fail in identifying causative genes underlying a given phenotype. Integrating GWAS and gene coexpression networks can help prioritize high-confidence candidate genes, as the expression profiles of trait-associated genes can be used to mine novel candidates. Here, we present *cageminer*, the first R package to prioritize candidate genes through the integration of GWAS and coexpression networks. Genes are considered high-confidence candidates if they pass all three filtering criteria implemented in *cageminer*, namely physical proximity to SNPs, coexpression with known trait-associated genes, and significant changes in expression levels in conditions of interest. Prioritized candidates can also be scored and ranked to select targets for experimental validation. By applying *cageminer* to a real data set, we demonstrate that it can effectively prioritize candidates, leading to >99% reductions in candidate gene lists.

**Availability and implementation:** The package is available at Bioconductor (http://bioconductor.org/packages/cageminer).

## 1 Introduction

Over the years, several genome-wide association studies (GWAS) have identified single-nucleotide polymorphisms (SNPs) associated with phenotypes of interest, such as agronomic traits in crops, production traits in livestock, and complex human disorders (Boudhrioua *et al.*, 2020; Maldonado Dos Santos *et al.*, 2019; Wu *et al.*, 2020; Buzanskas *et al.*, 2014). However, finding causative genes from SNPs remains a major bottleneck (Baxter, 2020). First, most GWAS-derived SNPs are located in non-coding portions of the genome, which can be regulatory regions very far from a causative gene (Peat *et al.*, 2020). Further, causative variants can be in strong linkage disequilibrium (LD) with non-causative ones, leading to large LD blocks with dozens of putative candidates (Michno *et al.*, 2020).

To address this issue, integrating GWAS with the vast amounts of RNA-seq data in public repositories has become a promising solution, particularly using gene coexpression network (GCN)-based approaches (Michno *et al.*, 2020; Yao *et al.*, 2020; Guo *et al.*, 2020). Currently, the only statistical framework that automates such integration is Camoco, a Python library that identifies sets of densely connected genes for a given sliding window relative to each SNP (Schaefer *et al.*, 2018). However, as sliding windows are expanded (*e.g.,* 50 kb), Camoco loses the ability to discover candidate genes because of background noise (Michno *et al.*, 2020). This is a major limitation, as SNPs can be up to 2 Mb away from the causative genes if they are in distal regions (Brodie *et al.*, 2016).

Here, we present *cageminer* (candidate gene miner), the first R/Bioconductor package that integrates GCNs and GWAS-derived SNPs to prioritize candidate genes associated with traits of interest. *cageminer* uses a guide gene-based approach to discover novel candidates that are coexpressed with known trait-associated genes and are significantly induced or repressed in conditions of interest. By relying on researchers’ prior knowledge, *cageminer* can identify high-confidence candidate genes even in megabase-scale genomic intervals. This package will be instrumental in helping researchers discover genes underlying important quantitative traits.

## 2 Implementation

*cageminer* is implemented as an R package, and all input and output objects belong to base R or common Bioconductor classes to ensure interoperability with other packages. Our algorithm requires three types of input data: i. SNP positions, which must be passed as GRanges or GRangesList objects (for single trait and multiple traits, respectively) (Lawrence *et al.*, 2013); ii. guide genes, either as a character vector or a data frame; and iii. gene coexpression network, which must be passed as a list as returned by the function *exp2gcn()* from the Bioconductor package *BioNERO* (Almeida-Silva and Venancio, 2021).

### 2.1 Algorithm description

*cageminer* identifies high-confidence candidate genes in three sequential steps (Fig. 1). In the first step, all genes within a sliding window (default: 2 Mb) relative to each SNP are selected as putative candidates. The default 2 Mb sliding window aims to minimize false-negative rates, as SNPs can be located in distal regions (Brodie *et al.*, 2016). If the 2 Mb window returns too many genes to start with, users can simulate different window sizes and visualize the number of genes in a line plot (Supplementary Text). Additionally, users can input a custom interval for each SNP (*e.g.,* based on linkage disequilibrium) by disabling the sliding window expansion.

**Fig. 1.**
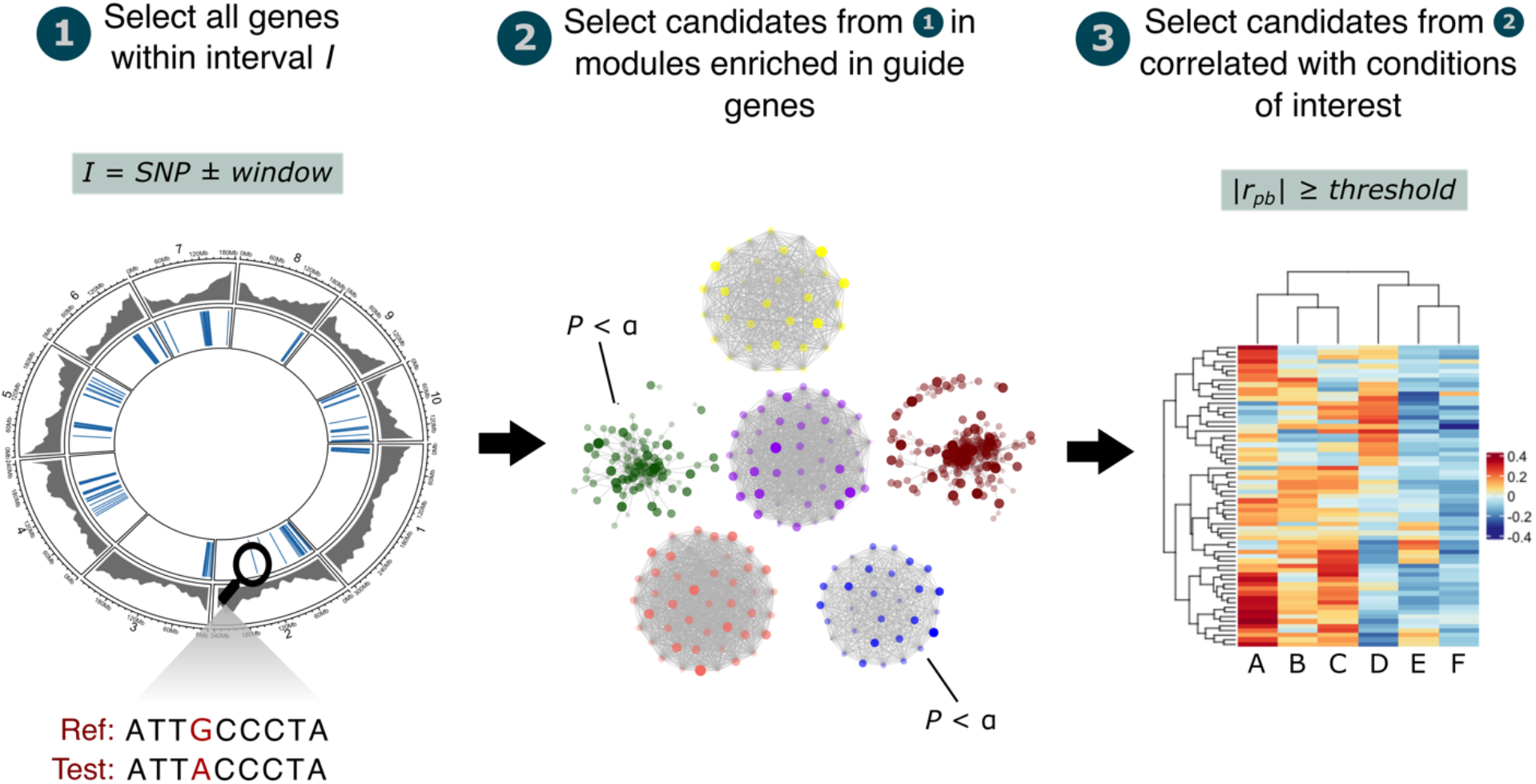
Summary of the *cageminer* algorithm. Candidate gene prioritization is performed in three sequential steps that can be run as a pipeline (recommended) or independently. The steps can be interpreted as different sources of evidence that candidates are causative genes. Thus, candidates that pass all three steps are considered high-confidence candidates.

For the second step of the algorithm, *cageminer* relies on the *module_enrichment()* function from the *BioNERO* package (Almeida-Silva and Venancio, 2021) to perform an enrichment analysis and find candidates from step 1 that co-occur in modules enriched in guide genes. Guides are genes known to be associated with the phenotype of interest, which can be passed as a single gene set in a character vector or as a 2-column data frame with gene IDs in the first column and gene classification (*e.g.,* Gene Ontology Terms or KEGG pathways) in the second column. In the latter case, *cageminer* will look for modules enriched in each class of guide genes rather than guides in general.

In the third step, the gene expression matrix used to infer the GCN is correlated to a binary matrix m_ij_ containing 1 if the sample *m* corresponds to the condition *j*, and 0 otherwise. This calculation, also known as gene significance, returns a point biserial correlation coefficient (r_pb_) (Langfelder and Horvath, 2008) that indicates if genes have significantly increased or decreased expression levels in a particular condition. Further, as genes can be negative regulators of the phenotype of interest, negative correlation coefficients are also treated as biologically meaningful. Thus, the absolute value of r_pb_ is considered to define a gene significance threshold, as well as Student asymptotic *P*-values for correlation significance (by default, r_pb_ ≥0.2 and *P* <0.05).

### 2.2 Gene scoring

To score the prioritized candidate genes and further select the top *n* genes for validation, genes can be scored with the formula below:

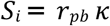

where

κ = 2 if the gene is a transcription factor

κ = 2 if the gene is a hub

κ = 3 if the gene is a hub and a transcription factor

κ = 1 if the gene is neither a hub nor a transcription factor

## 3 Application to a real dataset

A use case using RNA-seq on pepper (*Capsicum annuum*) response to Phytophthora root rot (Kim *et al.*, 2018), as well as GWAS SNPs associated with resistance to Phytophthora root rot (Siddique *et al.*, 2019) is available in the Supplementary Text. Pepper genes encoding transcription factors were downloaded from PlantTFDB 4.0 (Jin *et al.*, 2017), and plant defense-related genes (MapMan annotations) were obtained from PLAZA Dicots 3.0 (Proost *et al.*, 2015). From a list of 1265 putative candidates, *cageminer* identified 5 high-confidence candidate resistance genes (99.6% reduction). All candidates encode proteins related to known plant immunity-related processes (*e.g.*, immune signaling, oxidative stress, and lignan biosynthesis), supporting the effectiveness of the algorithm in finding biologically meaningful genes.

## 4 Conclusions

*cageminer* is the first R package to integrate GWAS-derived SNPs and gene coexpression networks to prioritize candidate genes involved in phenotypes of interest. This package will likely contribute to the advancement of population genomics and to the identification of genes for biotechnological applications.

## Acknowledgements

This work was supported by Fundação Carlos Chagas Filho de Amparo à Pesquisa do Estado do Rio de Janeiro (FAPERJ; grants E-26/203.309/2016 and E-26/203.014/2018), Coordenação de Aperfeiçoamento de Pessoal de Nível Superior - Brasil (CAPES; Finance Code 001), and Conselho Nacional de Desenvolvimento Científico e Tecnológico. The funding agencies had no role in the design of the study and collection, analysis, and interpretation of data and in writing.

