## Supplementary Text for "cageminer: an R/Bioconductor package to prioritize candidate genes by integrating GWAS and gene coexpression networks"

**4 August 2021**

### 1 Data description

The example data sets we will use here comprise RNA-seq data on pepper (*Capsicum annuum*) response to Phytophthora root rot (Kim et al. 2018), and GWAS-derived SNPs associated to resistance to Phytophthora root rot (Siddique et al. 2019). All genomic intervals are stored in GRanges objects, and expression data with sample metadata are stored in SummarizedExperiment objects. Genes encoding transcription factors were retrieved from PlantTFDB 4.0 (Jin et al. 2017), and plant defense-related genes (MapMan annotations) were retrieved from PLAZA Dicots 3.0 (Proost et al. 2015). Taking a glimpse at the data:

```
library(cageminer, quietly = TRUE)
##
## Registered S3 method overwritten by 'GGally':
##   method from
##   +.gg   ggplot2
set.seed(123) # for reproducibility

# SNP positions
data(snp_pos)
snp_pos
## GRanges object with 116 ranges and 0 metadata columns:
##       seqnames    ranges strand
##          <Rle> <IRanges>  <Rle>
##     2    Chr02 149068682      *
##     3    Chr03   5274098      *
##     4    Chr05  27703815      *
##     5    Chr05  27761792      *
##     6    Chr05  27807397      *
##   ...      ...       ...    ...
##   114    Chr12 230514706      *
##   115    Chr12 230579178      *
##   116    Chr12 230812962      *
##   117    Chr12 230887290      *
##   118    Chr12 231022812      *
##   -------
##   seqinfo: 8 sequences from an unspecified genome; no seqlengths

# Gene positions
data(gene_ranges)
gene_ranges
## GRanges object with 30242 ranges and 6 metadata columns:
##           seqnames              ranges strand |   source     type     score
##              <Rle>           <IRanges>  <Rle> | <factor> <factor> <numeric>
##       [1]    Chr01         63209-63880      - |  PGA1.55     gene        NA
##       [2]    Chr01       112298-112938      - |  PGA1.55     gene        NA
##       [3]    Chr01       117979-118392      + |  PGA1.55     gene        NA
##       [4]    Chr01       119464-119712      + |  PGA1.55     gene        NA
##       [5]    Chr01       119892-120101      + |  PGA1.55     gene        NA
##       ...      ...                 ...    ... .      ...      ...       ...
##   [30238]    Chr12 235631138-235631467      - |  PGA1.55     gene        NA
##   [30239]    Chr12 235642644-235645110      + |  PGA1.55     gene        NA
```

```
## [30240]    Chr12 235645483-235651927      - |  PGA1.55     gene        NA
## [30241]    Chr12 235652709-235655955      - |  PGA1.55     gene        NA
## [30242]    Chr12 235663655-235665276      - |  PGA1.55     gene        NA
##                phase          ID          Parent
##            <integer> <character> <CharacterList>
##       [1]      <NA>  CA01g00010
##       [2]      <NA>  CA01g00020
##       [3]      <NA>  CA01g00030
##       [4]      <NA>  CA01g00040
##       [5]      <NA>  CA01g00050
##       ...       ...         ...             ...
##   [30238]      <NA>  CA12g22890
##   [30239]      <NA>  CA12g22900
##   [30240]      <NA>  CA12g22910
##   [30241]      <NA>  CA12g22920
##   [30242]      <NA>  CA12g22930
##   -------
##   seqinfo: 12 sequences from an unspecified genome; no seqlengths

# Expression data in FPKM
data(pepper_se)
pepper_se
## class: SummarizedExperiment
## dim: 3892 45
## metadata(0):
## assays(1): ''
## rownames(3892): CA02g23440 CA02g05510 ... CA03g35110 CA02g12750
## rowData names(0):
## colnames(45): PL1 PL2 ... TMV-P0-3D TMV-P0-Up
## colData names(1): Condition

# Chromosome lengths
data(chr_length)
chr_length
## GRanges object with 12 ranges and 0 metadata columns:
##        seqnames      ranges strand
##           <Rle>   <IRanges>  <Rle>
##    [1]    Chr01 1-272704604      *
##    [2]    Chr02 1-171128871      *
##    [3]    Chr03 1-257900543      *
##    [4]    Chr04 1-222584275      *
##    [5]    Chr05 1-233468049      *
##    ...      ...         ...    ...
##    [8]    Chr08 1-145103255      *
##    [9]    Chr09 1-252779264      *
##   [10]    Chr10 1-233593809      *
##   [11]    Chr11 1-259726002      *
##   [12]    Chr12 1-235688241      *
##   -------
##   seqinfo: 12 sequences from an unspecified genome; no seqlengths
```

```
# Guide genes
data(guides)
head(guides)
##          Gene                              Description
## 1 CA10g07770                     response to stimulus
## 2 CA10g07770                       response to stress
## 3 CA10g07770            cellular response to stimulus
## 4 CA10g07770              cellular response to stress
## 6 CA10g07770 regulation of cellular response to stress
## 8 CA10g07770        regulation of response to stimulus

# Genes enconding TFs
data(tfs)
head(tfs)
##      Gene_ID Family
## 1 CA12g20650    RAV
## 2 CA00g00130   WRKY
## 3 CA00g00230   WRKY
## 4 CA00g00390    LBD
## 5 CA00g03050    NAC
## 6 CA00g07140 E2F/DP
```

### 2 Exploratory analysis

Before proceeding to the prioritization steps, it is important to explore the data to look for biologically relevant patterns. First, we can see if SNPs are evenly distributed across chromosomes or if they tend to co-occur in particular chromosomes.

```
plot_snp_distribution(snp_pos)
```

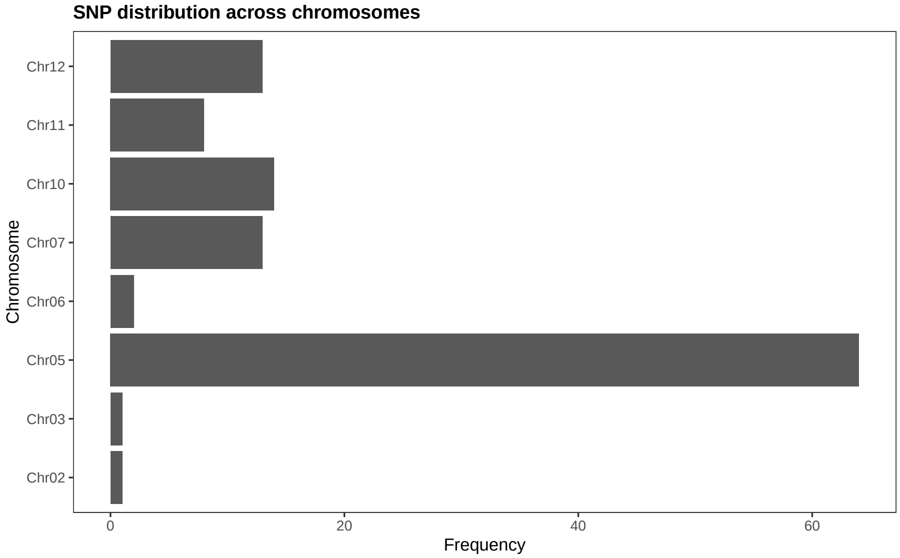

We can see that SNPs cluster in chromosome 05. Now, we can see if these SNPs in chromosome 05 are physically close to each other.

```
plot_snp_circos(chr_length, gene_ranges, snp_pos)
```

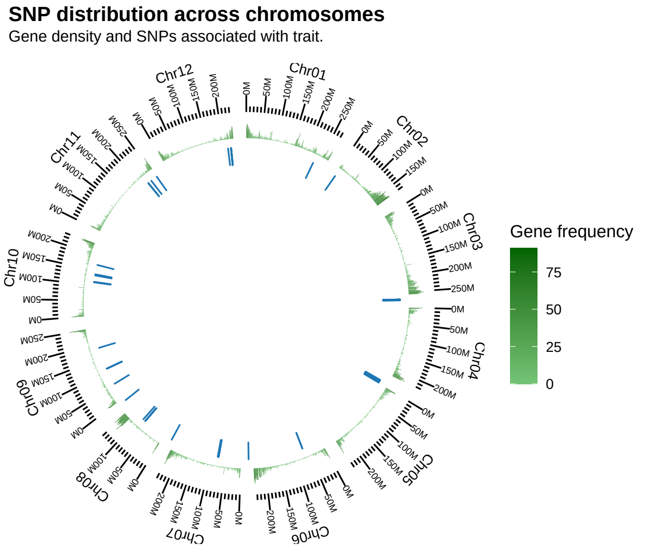

Indeed, they are very close to each other. Finally, we can simulate different sliding windows for the first step of the algorithm (see main text for details) to pick a custom interval.

```
simulate_windows(gene_ranges, snp_pos)
```

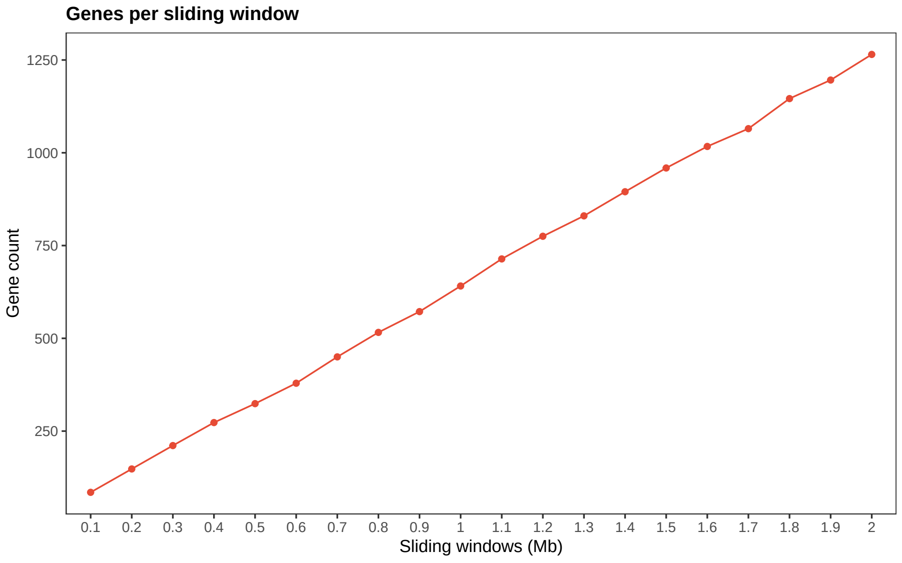

The plot shows that we can use the default sliding window (2 Mb), as it does not include too many genes. If information on linkage disequilibrium-based genomic intervals is available, we recommend using it.

### 3 Candidate gene prioritization

The three prioritization steps described in the paper can be applied with the functions `mine_step1()`, `mine_step2()`, and `mine_step3()`. Alternatively, the function `mine_candidates()` is a wrapper that combines the three `mine_*` functions to perform candidate gene prioritization in a single step. For the step 2, we will need to infer the gene coexpression network beforehand with the function `exp2gcn()` from the Bioconductor package BioNERO (Almeida-Silva and Venancio 2021).

```
# Apply step 1
step1 <- mine_step1(gene_ranges, snp_pos)
step1
## GRanges object with 1265 ranges and 6 metadata columns:
##          seqnames              ranges strand |   source     type     score
##             <Rle>           <IRanges>  <Rle> | <factor> <factor> <numeric>
##      [1]    Chr02 147076830-147083477      + |  PGA1.55     gene        NA
##      [2]    Chr02 147084450-147086637      - |  PGA1.55     gene        NA
##      [3]    Chr02 147099482-147104002      - |  PGA1.55     gene        NA
##      [4]    Chr02 147126373-147126537      + |  PGA1.55     gene        NA
##      [5]    Chr02 147129897-147132335      - |  PGA1.55     gene        NA
##      ...      ...                 ...    ... .      ...      ...       ...
##   [1261]    Chr12 232989761-232990947      - |  PGA1.55     gene        NA
##   [1262]    Chr12 232994658-232999784      + |  PGA1.55     gene        NA
##   [1263]    Chr12 233001307-233004705      + |  PGA1.55     gene        NA
##   [1264]    Chr12 233005539-233011740      - |  PGA1.55     gene        NA
##   [1265]    Chr12 233018159-233022142      - |  PGA1.55     gene        NA
##              phase          ID          Parent
##          <integer> <character> <CharacterList>
##      [1]      <NA>  CA02g16550
##      [2]      <NA>  CA02g16560
##      [3]      <NA>  CA02g16570
##      [4]      <NA>  CA02g16580
##      [5]      <NA>  CA02g16590
##      ...       ...         ...             ...
##   [1261]      <NA>  CA12g21190
##   [1262]      <NA>  CA12g21200
##   [1263]      <NA>  CA12g21210
##   [1264]      <NA>  CA12g21220
##   [1265]      <NA>  CA12g21230
##   -------
##   seqinfo: 12 sequences from an unspecified genome; no seqlengths

# Infer the GCN
sft <- BioNERO::SFT_fit(pepper_se, cor_method = "pearson")
## Warning: executing %dopar% sequentially: no parallel backend registered
##    Power SFT.R.sq   slope truncated.R.sq mean.k. median.k. max.k.
## 1      3 0.000902  0.0985          0.806   718.0    701.00 1060.0
## 2      4 0.039500 -0.4680          0.833   470.0    451.00  807.0
## 3      5 0.110000 -0.6540          0.851   322.0    301.00  639.0
## 4      6 0.269000 -0.9120          0.891   229.0    209.00  520.0
## 5      7 0.449000 -1.1200          0.920   168.0    149.00  432.0
```

```
## 6      8 0.598000 -1.2900          0.945   126.0    109.00  364.0
## 7      9 0.685000 -1.4300          0.949    96.8     81.00  311.0
## 8     10 0.744000 -1.5000          0.961    75.7     61.30  268.0
## 9     11 0.786000 -1.5800          0.964    60.2     47.00  233.0
## 10    12 0.817000 -1.6100          0.969    48.5     36.50  204.0
## 11    13 0.824000 -1.6600          0.966    39.5     28.80  180.0
## 12    14 0.831000 -1.6900          0.965    32.5     23.00  159.0
## 13    15 0.846000 -1.7000          0.972    27.1     18.30  142.0
## 14    16 0.859000 -1.7100          0.976    22.7     14.70  127.0
## 15    17 0.869000 -1.7200          0.981    19.2     11.90  115.0
## 16    18 0.877000 -1.7200          0.984    16.3      9.76  103.0
## 17    19 0.882000 -1.7200          0.986    14.0      7.97   93.7
## 18    20 0.889000 -1.7100          0.988    12.0      6.63   85.2
gcn <- BioNERO::exp2gcn(pepper_se, cor_method = "pearson", SFTpower = sft$power)
## ..connectivity..
## ..matrix multiplication (system BLAS)..
## ..normalization..
## ..done.

# Apply step 2
step2 <- mine_step2(pepper_se, gcn, guides, step1$ID)
## Enrichment analysis for module brown...
## Enrichment analysis for module cyan...
## Enrichment analysis for module darkgreen...
## Enrichment analysis for module darkmagenta...
## Enrichment analysis for module darkolivegreen...
## Enrichment analysis for module darkorange...
## Enrichment analysis for module darkorange2...
## Enrichment analysis for module darkred...
## Enrichment analysis for module darkturquoise...
## Enrichment analysis for module green...
## Enrichment analysis for module grey60...
## Enrichment analysis for module ivory...
## Enrichment analysis for module lightcyan...
## Enrichment analysis for module midnightblue...
## Enrichment analysis for module orange...
## Enrichment analysis for module orangered4...
## Enrichment analysis for module paleturquoise...
## Enrichment analysis for module pink...
## Enrichment analysis for module red...
## Enrichment analysis for module royalblue...
## Enrichment analysis for module salmon...
## Enrichment analysis for module steelblue...
## Enrichment analysis for module violet...
step2$candidates
##  [1] "CA10g08490" "CA03g01790" "CA10g12640" "CA12g21230" "CA10g02810"
##  [6] "CA03g01800" "CA02g17460" "CA10g02800" "CA03g03320" "CA05g14230"
## [11] "CA07g04010" "CA05g06480" "CA03g02720" "CA10g02630" "CA12g18010"
## [16] "CA07g04000" "CA02g16570" "CA02g17710" "CA10g02570" "CA05g15120"
## [21] "CA12g20980" "CA02g16830" "CA12g18440" "CA12g18400" "CA11g08940"
## [26] "CA10g02780" "CA12g19670" "CA07g12720" "CA03g01900" "CA12g07460"
```

```
## [31] "CA03g02360" "CA02g16620" "CA10g08420" "CA03g02960" "CA03g03010"
## [36] "CA07g12840" "CA05g15110" "CA02g16550" "CA05g14730" "CA02g16900"
## [41] "CA03g03310" "CA02g17030"

# Apply step 3
step3 <- mine_step3(pepper_se, candidates = step2$candidates,
                    sample_group = "PRR_stress")
```

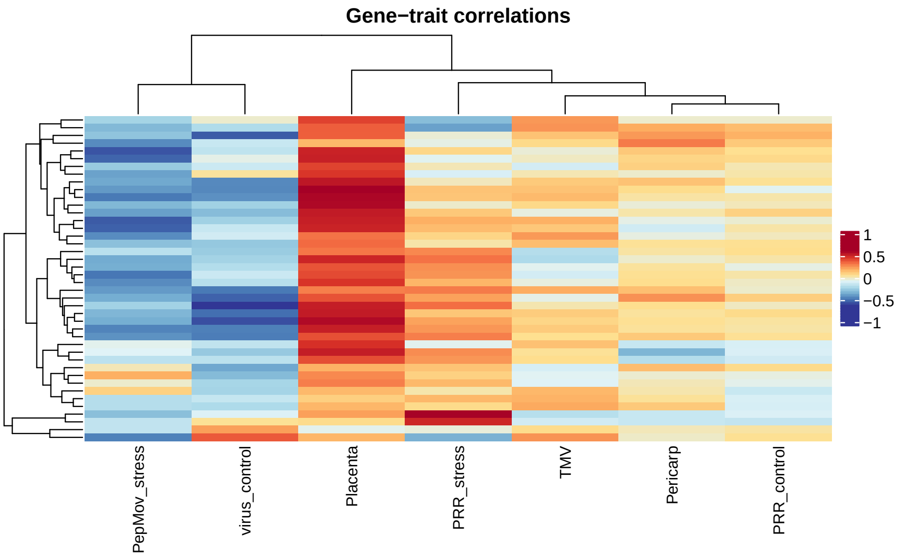

```
step3
##           gene      trait        cor       pvalue
## 264 CA12g18400 PRR_stress  0.5963394 1.540534e-05
## 243 CA11g08940 PRR_stress  0.4909160 6.171861e-04
## 201 CA10g02780 PRR_stress  0.3201048 3.205995e-02
## 19  CA02g16620 PRR_stress  0.3113993 3.732128e-02
## 33  CA02g16900 PRR_stress -0.3204388 3.187100e-02
## 180 CA07g12840 PRR_stress -0.3566806 1.617019e-02
## 110 CA03g03310 PRR_stress -0.3983772 6.720204e-03
```

### 4 Gene scoring

To conclude, we can score the prioritized candidates and rank them from highest to lowest score.

```
# Get hubs
hubs <- BioNERO::get_hubs_gcn(pepper_se, gcn)
scored_candidates <- score_genes(step3, hubs, tfs)
## Number of genes < 'pick_top'. Picking all genes.
scored_candidates
##           gene      trait        cor       pvalue     score
## 264 CA12g18400 PRR_stress  0.5963394 1.540534e-05  0.5963394
## 243 CA11g08940 PRR_stress  0.4909160 6.171861e-04  0.4909160
```

```
## 110 CA03g03310 PRR_stress -0.3983772 6.720204e-03 -0.3983772
## 180 CA07g12840 PRR_stress -0.3566806 1.617019e-02 -0.3566806
## 33  CA02g16900 PRR_stress -0.3204388 3.187100e-02 -0.3204388
## 201 CA10g02780 PRR_stress  0.3201048 3.205995e-02  0.3201048
## 19  CA02g16620 PRR_stress  0.3113993 3.732128e-02  0.3113993
```

Here, as none of the genes are hubs or TFs, their scores were represented by the $r_{pb}$ coefficients themselves.

### Session info

```
## R version 4.1.0 (2021-05-18)
## Platform: x86_64-apple-darwin17.0 (64-bit)
## Running under: macOS High Sierra 10.13.6
##
## Matrix products: default
## BLAS:   /Library/Frameworks/R.framework/Versions/4.1/Resources/lib/libRblas.dylib
## LAPACK: /Library/Frameworks/R.framework/Versions/4.1/Resources/lib/libRlapack.dylib
##
## locale:
## [1] en_US.UTF-8/en_US.UTF-8/en_US.UTF-8/C/en_US.UTF-8/en_US.UTF-8
##
## attached base packages:
## [1] stats     graphics  grDevices utils     datasets  methods   base
##
## other attached packages:
## [1] cageminer_0.99.6 BiocStyle_2.21.3
##
## loaded via a namespace (and not attached):
##   [1] utf8_1.2.2                  tidyselect_1.1.1
##   [3] RSQLite_2.2.7               AnnotationDbi_1.54.1
##   [5] htmlwidgets_1.5.3           grid_4.1.0
##   [7] BiocParallel_1.26.1         munsell_0.5.0
##   [9] codetools_0.2-18            preprocessCore_1.54.0
##  [11] statmod_1.4.36              colorspace_2.0-2
##  [13] OrganismDbi_1.34.0          filelock_1.0.2
##  [15] Biobase_2.52.0              knitr_1.33
##  [17] rstudioapi_0.13             stats4_4.1.0
##  [19] ggsignif_0.6.2              labeling_0.4.2
##  [21] MatrixGenerics_1.4.0        GenomeInfoDbData_1.2.6
##  [23] farver_2.1.0                bit64_4.0.5
##  [25] coda_0.19-4                 vctrs_0.3.8
##  [27] generics_0.1.0              xfun_0.24
##  [29] biovizBase_1.40.0           BiocFileCache_2.0.0
##  [31] fastcluster_1.2.3           markdown_1.1
##  [33] R6_2.5.0                    doParallel_1.0.16
##  [35] GenomeInfoDb_1.28.1         clue_0.3-59
##  [37] locfit_1.5-9.4              AnnotationFilter_1.16.0
##  [39] bitops_1.0-7                cachem_1.0.5
```

```
##  [41] reshape_0.8.8                DelayedArray_0.18.0        
##  [43] assertthat_0.2.1             BiocIO_1.2.0               
##  [45] networkD3_0.4                scales_1.1.1               
##  [47] nnet_7.3-16                  gtable_0.3.0               
##  [49] Cairo_1.5-12.2               sva_3.40.0                 
##  [51] WGCNA_1.70-3                 ggbio_1.40.0               
##  [53] ensembldb_2.16.3             rlang_0.4.11               
##  [55] BioNERO_1.1.1                genefilter_1.74.0          
##  [57] GlobalOptions_0.1.2          GENIE3_1.14.0              
##  [59] splines_4.1.0                rtracklayer_1.52.0         
##  [61] rstatix_0.7.0                lazyeval_0.2.2             
##  [63] impute_1.66.0                dichromat_2.0-0            
##  [65] broom_0.7.9                  checkmate_2.0.0            
##  [67] intergraph_2.0-2             BiocManager_1.30.16        
##  [69] yaml_2.2.1                   reshape2_1.4.4             
##  [71] abind_1.4-5                  GenomicFeatures_1.44.0     
##  [73] ggnetwork_0.5.10             backports_1.2.1            
##  [75] Hmisc_4.5-0                  gridtext_0.1.4             
##  [77] RBGL_1.68.0                  tools_4.1.0                
##  [79] bookdown_0.22                statnet.common_4.5.0       
##  [81] ggplot2_3.3.5                ellipsis_0.3.2             
##  [83] RColorBrewer_1.1-2           BiocGenerics_0.38.0        
##  [85] dynamicTreeCut_1.63-1        Rcpp_1.0.7                 
##  [87] plyr_1.8.6                   base64enc_0.1-3            
##  [89] progress_1.2.2               zlibbioc_1.38.0            
##  [91] purrr_0.3.4                  RCurl_1.98-1.3             
##  [93] prettyunits_1.1.1            ggpubr_0.4.0               
##  [95] rpart_4.1-15                 GetoptLong_1.0.5           
##  [97] cowplot_1.1.1                S4Vectors_0.30.0           
##  [99] SummarizedExperiment_1.22.0 haven_2.4.1                
## [101] cluster_2.1.2                magrittr_2.0.1             
## [103] magick_2.7.2                 data.table_1.14.0          
## [105] openxlsx_4.2.4               circlize_0.4.13            
## [107] ProtGenerics_1.24.0          ggnewscale_0.4.5           
## [109] matrixStats_0.60.0           hms_1.1.0                  
## [111] evaluate_0.14                xtable_1.8-4               
## [113] minet_3.50.0                 RhpcBLASctl_0.20-137       
## [115] XML_3.99-0.6                 rio_0.5.27                 
## [117] jpeg_0.1-9                   readxl_1.3.1               
## [119] IRanges_2.26.0               gridExtra_2.3              
## [121] shape_1.4.6                  compiler_4.1.0             
## [123] biomaRt_2.48.2               tibble_3.1.3               
## [125] crayon_1.4.1                 htmltools_0.5.1.1          
## [127] mgcv_1.8-36                  Formula_1.2-4              
## [129] ggtext_0.1.1                 tidyr_1.1.3                
## [131] geneplotter_1.70.0           DBI_1.1.1                  
## [133] dbplyr_2.1.1                 ComplexHeatmap_2.8.0       
## [135] rappdirs_0.3.3               Matrix_1.3-4               
## [137] car_3.0-11                   parallel_4.1.0             
## [139] igraph_1.2.6                 GenomicRanges_1.44.0       
## [141] forcats_0.5.1                pkgconfig_2.0.3
```

```
## [143] GenomicAlignments_1.28.0    foreign_0.8-81
## [145] xml2_1.3.2                  foreach_1.5.1
## [147] annotate_1.70.0             XVector_0.32.0
## [149] VariantAnnotation_1.38.0    stringr_1.4.0
## [151] digest_0.6.27               graph_1.70.0
## [153] NetRep_1.2.4                Biostrings_2.60.1
## [155] rmarkdown_2.9.5             cellranger_1.1.0
## [157] htmlTable_2.2.1             edgeR_3.34.0
## [159] restfulr_0.0.13             curl_4.3.2
## [161] Rsamtools_2.8.0             rjson_0.2.20
## [163] lifecycle_1.0.0             nlme_3.1-152
## [165] carData_3.0-4               network_1.17.1
## [167] BSgenome_1.60.0             limma_3.48.1
## [169] fansi_0.5.0                 pillar_1.6.1
## [171] lattice_0.20-44             GGally_2.1.2
## [173] KEGGREST_1.32.0             fastmap_1.1.0
## [175] httr_1.4.2                  survival_3.2-11
## [177] GO.db_3.13.0                glue_1.4.2
## [179] zip_2.2.0                   png_0.1-7
## [181] iterators_1.0.13            bit_4.0.4
## [183] stringi_1.7.3               blob_1.2.2
## [185] DESeq2_1.32.0               latticeExtra_0.6-29
## [187] memoise_2.0.0               dplyr_1.0.7
```
